# On the effects of selection and mutation on species tree inference

**DOI:** 10.1101/2020.09.08.288183

**Authors:** Matthew Wascher, Laura S. Kubatko

**Affiliations:** Ohio State University; The Ohio State University

## Abstract

A common question that arises when inferring species-level phylogenies from genome-scale data is whether selection acting on certain parts of the genome could create a bias in the inferred phylogeny. While most methods for species tree inference currently assume the multispecies coalescent (MSC), all methods that we are aware of utilize only the neutral coalescent process. If selection is in fact present, failure to adequately model it could introduce substantial bias. We work toward rigorously addressing this question using mathematical theory by deriving a version of the coalescent including selection and mutation as a limiting approximation of the Wright-Fisher model with selection and mutation, and showing that it can be used to closely approximate the distribution of coalescent times in the presence of selection and mutation. We confirm the adequacy of the approximation with a simulation study, and discuss its implications for species tree inference. Our results show that in a general class containing many cases of interest, selection has only a small impact on the coalescent process, and ignoring selection when it is present does not have a substantial negative impact on inference of the species tree topology.

## 1 Introduction

A common question that arises when inferring species-level phylogenies from genome-scale data is whether selection acting on certain parts of the genome could create a bias in the inferred phylogeny. While most methods for species tree inference currently assume the multispecies coalescent (MSC), all methods that we are aware of utilize only the neutral coalescent process. If selection is in fact present, these methods then use a misspecified model, which, in many estimation problems, introduces substantial bias. We work toward rigorously addressing this question using mathematical theory to show that in a general class containing many cases of interest, ignoring selection when it is present does not have a substantial negative impact on species tree inference. We confirm our results empirically using simulation.

The primary complication involved in moving from the neutral coalescent model to the coalescent with selection is that the property of *exhangeability* among lineages no longer applies, as certain allele types become differentially likely to survive to the next generation. The loss of exchangeability poses significant computational challenges that have impeded the incorporation of the coalescent with selection in methods for species tree estimation. Although the coalescent with selection has been well-studied (see, e.g., Wakeley (2009)), these computational difficulties are likely a major reason that methods for species tree inference assume the neutral coalescent.

Within the field of population genetics, the modern literature concerning the role of selection at the population level begins with Kaplan et al. (1988), who examine the population dynamics, follow the approach set out by Kimura (1955) in finding a continuous diffusion process that approximates these dynamics, and then study the properties of this diffusion process. A great deal of subsequent literature follows this diffusion approximation strategy, and quite a lot is known about these processes, including convergence results in a number of cases involving selection, mutation, and recombination that allow limiting expressions for coalescent times to be derived in closed form. For an introduction to the diffusion approach in population genetics and a review of results in this area, we direct the reader to Wakeley (2009), Barton and Etheridge (2019), and Svirezhev and Passekov (1990).

However, we aim here to study the coalescent with selection from the perspective of phylogenetic inference, which necessitates computing tree probabilities. From this perspective, diffusion approximations pose several challenges. First, limiting expressions for the coalescent times will generally depending on having information about at least some subset of the mutation rate, selective advantage, and allele types of individuals at all times – information that is not typically available in the phylogenetic inference setting. Second, even when an expression for the coalescent times is known, the simplicity of computing tree probabilities by using symmetry and counting is lost. It quickly becomes infeasible to compute tree probabilities as trees grow in size.

Previous work has, to some extent, considered the effect of selection on genome-scale phylogenetic inference. For example, Siepel (2009) discusses the role of background selection in forcing local reductions in effective population size along a chromosome, which has the effect of increasing the rate of coalescence in the region linked to the selected sites, resulting in less incomplete lineage sorting (ILS). Siepel (2009) further noted that the work of McVicker et al. (2009) provides empirical support for this process via examination of the local genealogies in neutral regions that are linked to constrained genes in hominoid genomes. Others (see, e.g., Rannala and Yang (2003); Zhu and Yang (2012); Edwards (2009); Edwards et al. (2016)) have also argued that certain types of selection result in reduced levels of ILS and are thus not problematic for genome-scale inference. In particular, Edwards (2009) considered the effects of various types of selective pressure on gene phylogenies, and pointed out that balancing selection tends to produce phylogenies that respect species-level relationships to an even greater extent than neutral selection. An additional argument in favor of the robustness of species tree inference methods is the fact that a relatively small percentage of the genome is generally believed to be under selection, and thus the signal coming from the neutral loci should be able to overwhelm the signal from the few loci under selection when genome-scale data is being analyzed.

Several studies, on the other hand, have highlighted the potentially important role of selection in phylogenomic inference. One argument about the importance of selection is that it is known that selection may affect gene tree inference Castoe et al. (2009); McVicker et al. (2009), thus suggesting that data collected from loci under selection be discarded prior to species tree inference (Rannala and Yang, 2003; Yang and Rannala, 2010; Zhang et al., 2011; Springer and Gatesy, 2016). However, this leads to the complicated task of identifying such loci (Adam et al., 2018). Furthermore, some studies argue that the proportion of the genome affected by some type of selection may be much larger than often assumed (Hahn, 2008; McVicker et al., 2009; Scally et al., 2012; Corbett-Detig et al., 2015). Empirical evidence to this effect has been noted by Silva et al. (2015), who found that between 16% and 33% of genes were under positive selection in rust fungi, and Tong et al. (2017), who found that phylogenies corresponding to genes under negative selection in insects showed different patterns than those subject only to drift.

While the work described above focuses primarily on the effect of selection for single gene phylogenies across the genome, as well as the consideration of what proportion of the genome is likely to be subject to selective pressure, little formal work has explicitly addressed the effect of selection on inference of the species-level phylogeny and on the estimation of associated parameters. A notable exception is the work of Adam et al. (2018), who use simulation to examine the impact of selection of varying strength within one species in a three-tip species tree on both the accuracy of the resulting estimate of the species tree and on parameters such as effective population size and divergence time. By examining the distribution of simulated gene trees under selection, Adam et al. (2018) confirmed the prediction that positive selection leads to an increased probability of monophyly within the species under selection, increasing the chance that the gene tree matches the species tree, even at shallow divergences, though it was possible to perturb the gene tree distribution under a scenario of weak selection at deep divergences. In terms of estimation of the species tree topology, the method examined by Adam et al. (2018), BPP (Rannala and Yang, 2017; Flouris et al., 2018), was generally robust to selection regardless of its strength, though the presence of strong selection that affected a large proportion of loci did negatively affect estimation accuracy. However, selection had the effect of biasing species divergence time estimates to be too large. Overall, Adam et al. (2018) conclude that, while selection has the largest effect when sample sizes within species are large, species are recently diverged, and strong selection is present at multiple loci, the accuracy of species tree estimates was largely unaffected by the conditions that are most common in empirical studies.

We aim to examine this question from a more theoretical perspective. We begin by studying the Markov chain governing the underlying population dynamics, following Kaplan et al. (1988)) Then, rather than use a diffusion approximation, we show how a discrete-time approximation can be used to see that in the limit as the population size grows, for fixed selective advantage and mutation rate, the coalescent with selection is equivalent to the classical coalescent with time rescaled, although the time rescaling changes each time a coalescent event occurs. We then discuss when this limiting approximation is practically valid and discuss the consequences for phylogenetic inference in various cases. Our results generally agree with what was empirically observed by Adam et al. (2018).

## 2 Results

### 2.1 An approximation for the rate of coalescence with selection

In this section, we provide an overview of our approximation for the rate of coalescence under the classical coalescent model with selection and mutation. Full details of the derivation can be found in the Materials and Methods. In the next section, we apply our approximation to the problem of inferring a species-level phylogeny from phylogenomic data.

#### 2.1.1 The coalescent with selection

We consider the standard definition of the Wright-Fisher model with selection, details of which can be found in Wakeley (2009). This model assumes a fixed population size, *N*, with two types of individuals, type *A* and type *a*. Additionally we will assume that type *A* individuals have reproductive advantage *s >* 1, meaning that type *A* individuals are *s* times more likely to reproduce than type *a* individuals. Finally, we assume a mutation rate *p*_*m*_. Mutant offspring will be of the opposite type from their parents, while non-mutant offspring will be of the same type as their parent.

The model then proceeds analogously to the classical Wright-Fisher model, except that we sample the next generation from the current according to the rules generated by *s* and *p*_*m*_ above. This can be done as follows. Suppose that there are *n*_1_ type *A* individuals and *n*_2_ type *a* individuals, where *n*_1_ + *n*_2_ = *N*, in the current generation. To generate the *i*^*th*^ individual of the next generation, first generate a uniform random variable *U*_*i*_ ∼ Uniform[0, *sn*_1_ + *n*_2_], and select individual ⌈*U*_*i*_*/s*⌉ as the parent if *U*_*i*_ ≤ *sn*_1_ and individual ⌈*U*_*i*_ − *sn*_1_ + *n*_1_⌉ as the parent if *U*_*i*_ *> sn*_1_. Then flip a biased coin with probability of success *p*_*m*_. On a success, assign individual *i* the type opposite of its parent, otherwise assign individual *i* the type of its parent. Repeat this process independently for *i* = 1 … *N* to generate the *N* individuals of the next generation.

Our first goal is to understand how the proportion of individuals in the population with the advantageous allele *A* changes from generation to generation. We thus define the Markov chain 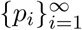 to track the proportion of type *A* individuals in the *i*^*th*^ generation. The transition probabilities of this Markov chain are given in the following lemma.

##### Lemma 2.1

(Genetic drift). *Let s be the fitness advantage of A, and let p*_*m*_ *be the mutation rate. Suppose the initial population has size N and proportion of A individuals p*_0_. *Then* 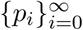 *is a Markov chain with transition probabilities given by*

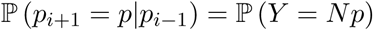

*where*

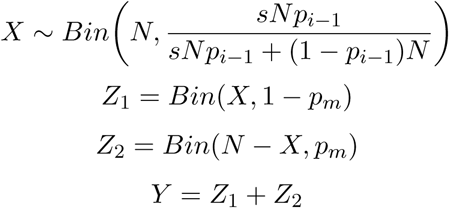

In the lemma above, *X* is the number of type *A* individuals chosen to reproduce in generation *i, Z*_1_ is the number of these that do not undergo mutation (and thus retain type *A*), and *Z*_2_ is the number of *a* individuals that mutate to type *A*. Since the number of *A* individuals in the *i*^*th*^ generation is just the sum two Binomially-distributed random variables (*Z*_1_ and *Z*_2_), the transition probabilities of the Markov chain *{p*_*i*_*}* are the sum of two Binomial probabilities. This leads to convenient properties, summarized by Lemmas (4.1) - (4.5) in the Materials and Methods, that are useful in determining the large-sample behavior of the Markov chain, as discussed in the following sections.

#### 2.1.2 Dynamics of the Markov chain

Our next goal is to understand the behavior of the Markov chain 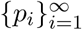, and in particular how it is affected by the strength of selection and the mutation rate. To study the behavior of the chain, we proceed in two steps. In this section, we show that *{p*_*i*_*}* quickly concentrates around a particular value *p*, which is a function of *s* and *p*_*m*_, and remains near *p* for a long time when *N* is sufficiently large. In the next section, we use this result to show that the rate at which lineages in a sample of size *M* coalesce is a rescaled version of the rate in the standard coalescent, where the rescaling depends on *p*. This allows a straightforward approach to examining the effect of selection and mutation on the distribution of gene tree topologies and branch lengths, and thus on coalescent-based species tree inference in the presence of selection and mutation.

We provide an outline of the methods used to derived these results. Full details and proofs are given in the Materials and Methods. To begin, we define the value *p* around which the Markov chain concentrates:

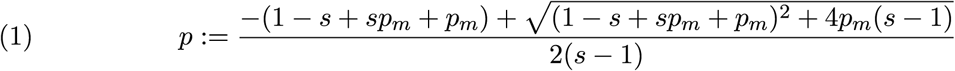

which is in [0, 1] when *s >*1 and *p*_*m*_ ∈ [0, 1]. We next show that if *p*_*t*_ is close to *p*, then it is very likely that *p*_*w*_ is also close to *p* for all *w > t* such that *w* − *t < N* ^*k*^ where we may choose *k* ≥ 2 when *N* is sufficiently large. We state this formally in the following theorem.

##### Theorem 2.2

*Choose any k* ∈ ℕ. *Consider the Markov chain* 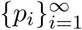 *with p*_0_ ∈ [0, 1] *and a population size of N*. *Then for δ, ϵ >* 0 *and sufficiently small, there exists N* ^∗^ *such that for N > N* ^∗^ *and all N* ^1*/k*^ ≤ *i* ≤ *N* ^*k*^,

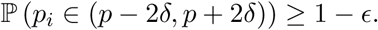

To show this, we first pick *N* ^∗^ large enough that 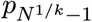 ∈ (*p* − *δ, p*+*δ*) with high probability. That this can be done is a consequence of Lemma 2.3 given below. We then consider the possible ways that *p*_*i*_ could leave the interval (*p* − 2*δ, p* + 2*δ*) given *p*_*i*−1_ is in (*p* − 2*δ, p* + 2*δ*). If in fact *p*_*i*−1_ ∈ (*p* − *δ, p* + *δ*), then in order for *p*_*i*_ to leave (*p* − 2*δ, p* + 2*δ*), the Markov chain must make a jump of size at least *δ* in one generation. Using properties of the Binomial transition probabilities given in Lemma 2.1, we show that the probability that this happens goes to 0 as *N* becomes large. If *p*_*i*−1_ ∈ (*p* − 2*δ, p* + 2*δ*) but not (*p* − *δ, p* + *δ*), then we apply Lemma 2.3 again to show that with high probability *{p*_*i*_*}* quickly reaches (*p* − *δ, p* + *δ*) again without leaving (*p* − 2*δ, p* + 2*δ*). Furthermore, in both cases the probability that *p*_*i*_ leaves (*p* − 2*δ, p* + 2*δ*) given *p*_*i*−1_ ∈ (*p* − *δ, p* + *δ*) is sufficiently small that we can apply a union bound over *N* ^*k*^ generations and still observe that the probability goes to 0 as *N* becomes large. Thus, the chain *{p*_*i*_*}* remains in the desired interval with probability as high as desired when the population size is sufficiently large. The formal proof is given in Materials and Methods.

Theorem 2.2 shows that once *{p*_*i*_*}* is close to *p*, it stays close to *p* for a long time. The utility of this result thus depends on the speed with which *{p*_*i*_*}* gets close to *p*. Our next result shows that this happens quickly.

##### Lemma 2.3

*For large N and any p*_0_ ∈ [0, 1] *\* (*p* − *δ, p* + *δ*), *with high probability the process {p*_*i*_*} moves monotonically toward the interval* (*p* − *δ, p* + *δ*) *and reaches it in at most p/C steps for some constant C >* 0 *that does not depend on N*.

In the next section, we show how these results can be used to rescale the rate of coalescence to account for the effects of selection and mutation.

#### 2.1.3 Scaling the rate of coalescence

Our previous results show that it is very likely that *{p*_*i*_*}* quickly gets close to *p* and stays there for at least *N* ^*k*^ generations for some *k* ≥ 2 that we may choose. As in the standard coalescent without selection, coalescent events happen at rate at least *O*(1*/N*). When *N* is large it is very unlikely that *{p*_*i*_*}* will deviate far from *p* before we see all lineages coalesce, assuming that the number of lineages is small relative to the population size and that we choose the value of *k* in Theorem 2.2 appropriately. This suggests that asymptotically the coalescent process behaves as if *p*_*i*_ = *p* for all *i*, and so this should be a reasonable approximation when *N* is large.

Accordingly, we now examine the coalescent process under the assumption that *p*_*i*_ = *p* for all *i*. Our goal is to show that when we consider time in appropriately scaled “coalescent units,” the waiting time until the next coalescent event is asymptotically an exponential random variable with some rate *λ* that will depend on *s, p*_*m*_, and the number of lineages *M*. Note that in the standard coalescent, the rate depends only on *M*.

We first need to compute the probability of a coalescent event in the previous generation. Note that this probability depends on how many lineages had type *A* versus type *a* parents, since two lineages cannot coalesce if they had different parent types. Suppose we have *h*^∗^ type *A* lineages and *M* − *h*^∗^ type *a* lineages, with *M* total lineages sampled from a population of size *N*. Then we can see that for a each lineage *L*_*type*_, we have that depending on its type

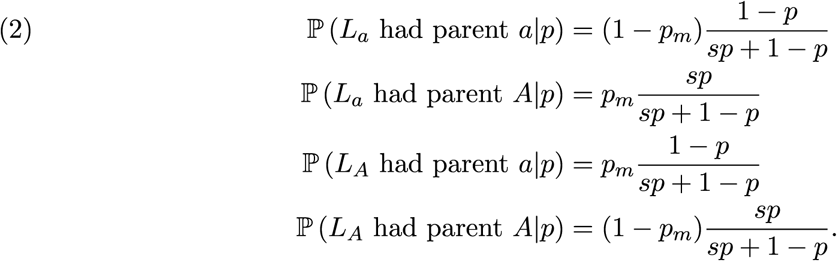

Suppose now that we have *h* lineages with type *A* parents and *M* − *h* lineages with type *a* parents. Exactly one coalescent event happens if we have either one coalescent event among the *h* type *A* parent lineages or one coalescent event among the *M* − *h* type *a* parent lineages but not both. Each of these follows the standard coalescent independently with *h* lineages and population size *Np* and *M* − *h* lineages and population size *N* (1 − *p*), respectively. Using the same argument as in the standard coalescent, we see that the probabilities of coalescence in the immediately previous generation are given by:

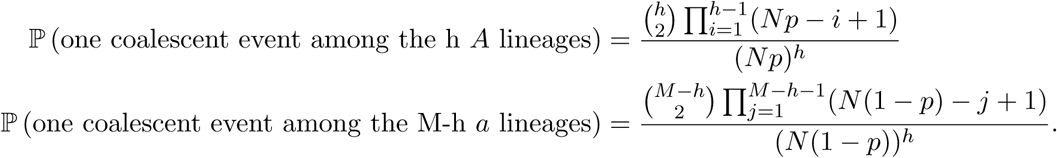

Ignoring the possibility of more than one coalescent event in the previous generation, as is assumed in the standard coalescent, the probability of observing a coalescent event in the immediately previous generation is

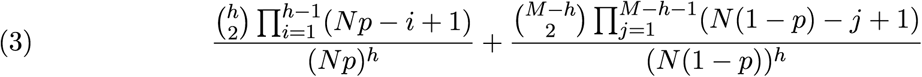

We would now like to treat the number of generations to the coalescent event as a geometric random variable with success probability given by the previous expression. However, unlike in the standard coalescent, the probability of success for this geometric random variable varies from generation to generation because it depends on the value of *h*. To deal with this, we use an argument based on coupled Markov chains to show that the Markov chain that tracks *h* mixes quickly when *M << N*. We then use the stationary distribution of this chain, denoted *π*(*h*), to approximate the proportion of time spent in each state *h*. Details can be found in the Materials and Methods.

Under these assumptions and counting time in units of *N* generations, let *T* ^*N*^ be the time until the coalescent event from *M* lineages to *M* − 1 lineages. In the Materials and Methods, we use the expression given in Equation 3 to show that when *N* is large, *T* ^*N*^ is approximately exponentially distributed with rate parameter

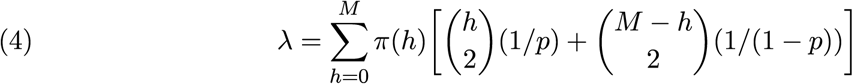

Although this expression looks simple, we note that *π*(*h*) is in fact a function of *M, s*, and *p*_*m*_ as well as *h*, which depends on these parameters in a complex way that involves matrix inversion. Thus, while it is not difficult to compute *λ* for given values of *M, s*, and *p*_*m*_, it is difficult to study the expression for *λ* theoretically.

Nonetheless, this approximation allows us to study the effect of selection and mutation on species-level phylogenomic inference by using standard calculations with the usual rate of coalescence, 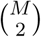, replaced by *λ*. We note that as *p* approaches 0 or 1 (corresponding to the case of no selection), *λ* approaches the value for the standard coalescent. After verifying the adequacy of this approximation, we look at how *λ* varies from the standard coalescent rate for different choices of *s, p*_*m*_, and *M*, and we examine the effect on the resulting distribution of gene trees.

### 2.2 Application to species tree inference

In this section, we use simulation to demonstrate that the approximation derived in the previous section is adequate for studying the distribution of times of coalescent events, which enables application of the results to the study of species tree inference. We then show that while the rate of coalescence is increased in the presence of selection, the effect is small under reasonable conditions, and thus that the distribution of gene trees deviates little from that expected in the absence of selection. The implications of these findings are explored in the Discussion.

#### 2.2.1 The approximation is adequate for studying coalescent-time distributions

To examine the adequacy of the approximation derived in the previous section, we simulated 1000 coalescent times when *M* = 4 and *N* = 200000 and when *M* = 6 and *N* = 500000 for all combinations of *s* ∈ *{*1.0002, 2*}* and *p*_*m*_ ∈ *{*0.0001, 0.1*}*. We compared the distributions of both *h* (the number of lineages of type *A*) and the time to the first coalescent event under our approximation to those simulated from the coalescent process with mutation and selection as described above. A detailed description of our simulation algorithm can be found in Materials and Methods and our code can be found at https://github.com/mwwascher/selection.

Figure 1 shows the simulated distribution of *h* and coalescent times when *s* = 2 and *p*_*m*_ = 0.1 and when *s* = 1.0002 and *p*_*m*_ = 0.0001 for the cases when *M* = 6 and *M* = 4. Similar plots for the other values of *s* and *p*_*m*_ can be found in the Supplemental Information. Although *p*_*m*_ = 0.1 is an unrealistically large mutation rate, we include this case because we expect to observe fast convergence even for a small population size if our approximation is valid, which is confirmed by Figure 1 (a)-(d). Alternatively, we expect convergence to be the slowest when both *s* − 1 and *p*_*m*_ are small, and thus the setting in Figure 1 (e)-(h) represents a more difficult condition. We note that even in this difficult setting, the simulated distribution is well-approximated by our method. Overall, we see good agreement between the coalescent times predicted by our approximation and those simulated, suggesting that our approximation will be useful in examining the impact of selection on species-level phylogenomic inference.

**Figure 1:**
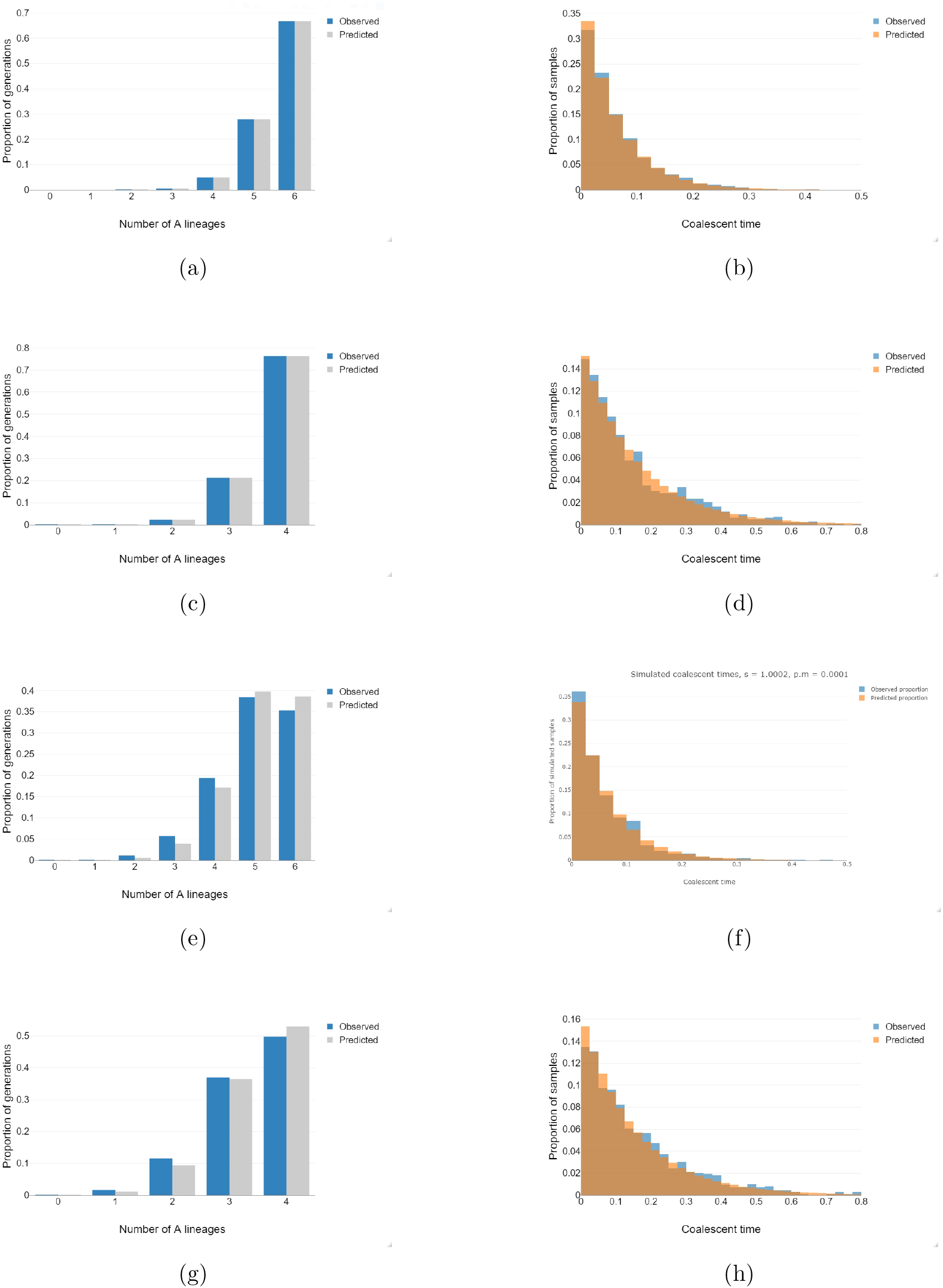
Comparison of the expected and approximated distributions of *h*, the number of *A* lineages, (left column) and the time to the first coalescent event (right column) for various choices of *M, s*, and *p*_*m*_. (a, b) *M* = 6, *s* = 2, *p*_*m*_ = 0.1; (c, d) *M* = 4, *s* = 2, *p*_*m*_ = 0.1; (e, f) *M* = 6, *s* = 1.0002, *p*_*m*_ = 0.0001; (g, h) *M* = 4, *s*= 1.0002, *p*_*m*_ = 0.0001.

#### 2.2.2 Mutation and selection have small effects on the rate of coalescence

A natural question that arises is how the value of *λ* predicted by the model with selection compares to *λ* when selection is not present. Figure 2 shows the value *λ* as a function of *s* and *p*_*m*_ when *M* = 4 (a) and when *M* = 6 (b). Noting that in the absence of selection the coalescent rates are 6 and 15 for *M* = 4 and *M* = 6, respectively, we see that selection increases the rate of coalescence by only a small amount, at most 10-15%. Additionally, we note that the rate is highest when *s* − 1 and *p*_*m*_ are comparable. This is intuitive, as when *s* − 1 is much larger than *p*_*m*_, selection overwhelms mutation and almost all of the population is type *A* in every generation, while when *s* − 1 is much smaller than *p*_*m*_, mutation overwhelms selection and type *A* individuals mutate before they can accumulate selective advantage over time. These results match what has been observed by others (e.g., Siepel (2009); Edwards (2009); Adam et al. (2018)), in that selection increases the rate of coalescence which has the effect of reducing the occurrence of ILS, making it more likely that the gene tree will match the species tree. However, our results suggest that this effect is small. We study this in further detail in the next section.

**Figure 2:**
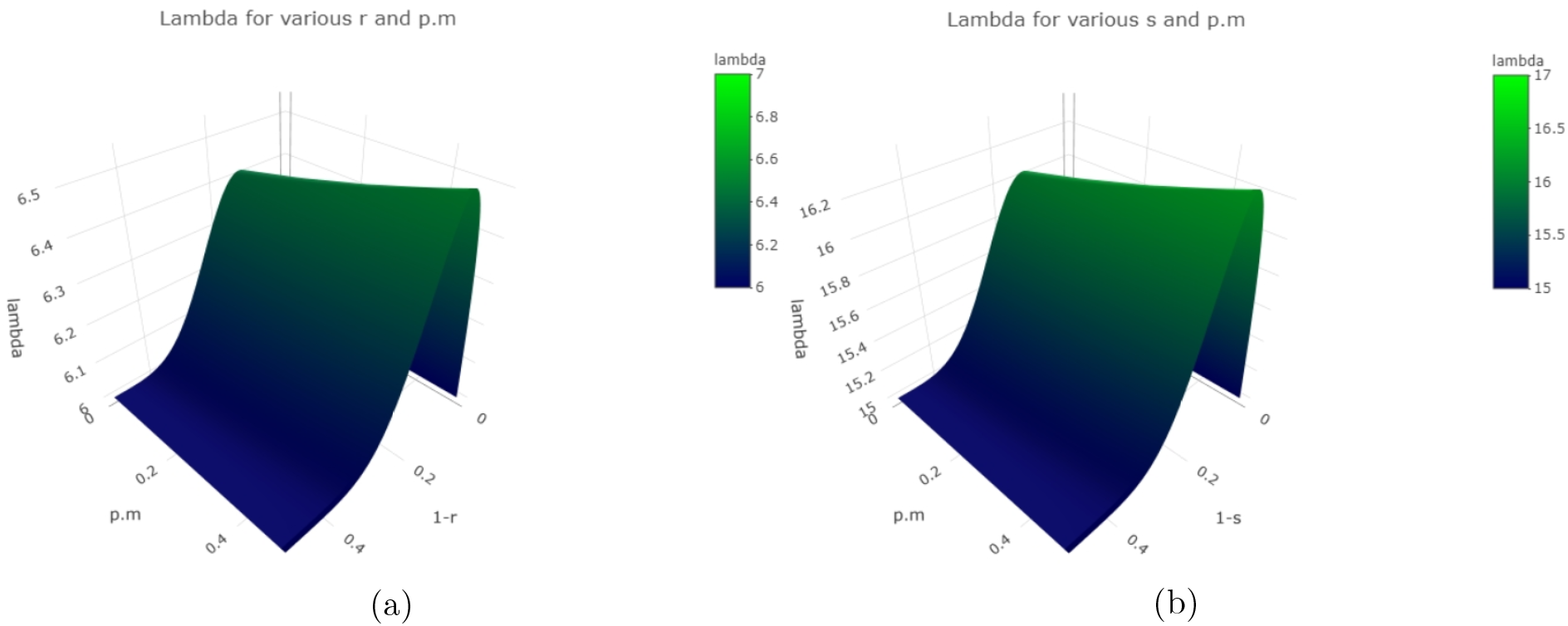
The rate of coalescence, *λ*, as a function of *s* and *p*_*m*_ for *M* = 4 (a) and for *M* = 6 (b).

#### 2.2.3 Mutation and selection have a small effect on the distribution of gene trees

In order to understand the impact of selection on phylogenetic inference, we need to understand how it affects the probabilities of different gene tree topologies given a species tree. As we saw in Section 2.2.2, selection increases the rate of coalescence. Intuitively, faster coalescence will lead to less ILS and make it more likely that we observe gene trees with topologies matching that of the species tree.

To investigate further, we simulated 10,000 gene trees from 4-taxon species trees with both the symmetric and the asymmetric topology with varying branch lengths for the neutral case and for the coalescent under selection for all combinations of *s* ∈ *{*1.0002, 2*}* and *p*_*m*_ ∈ *{*0.0001, 0.1*}*. Note: This will be replaced with calculation rather than simulation. Figure 3 shows the distribution of gene trees for various choices of the parameters for the symmetric species tree show in panel (a). A corresponding figure for the asymmetric species tree is given in the Supplemental Information. In panels (b) - (f), we see that the gene tree probabilities for the neutral coalescent (blue bars) are very close to those for various choices of *s* and *p*_*m*_, confirming that selection has little effect on the distribution of gene trees. In addition, we see in panel (b) that the probability of the gene tree that matches the species tree is higher for all of the scenarios in which we have selection.

**Figure 3:**
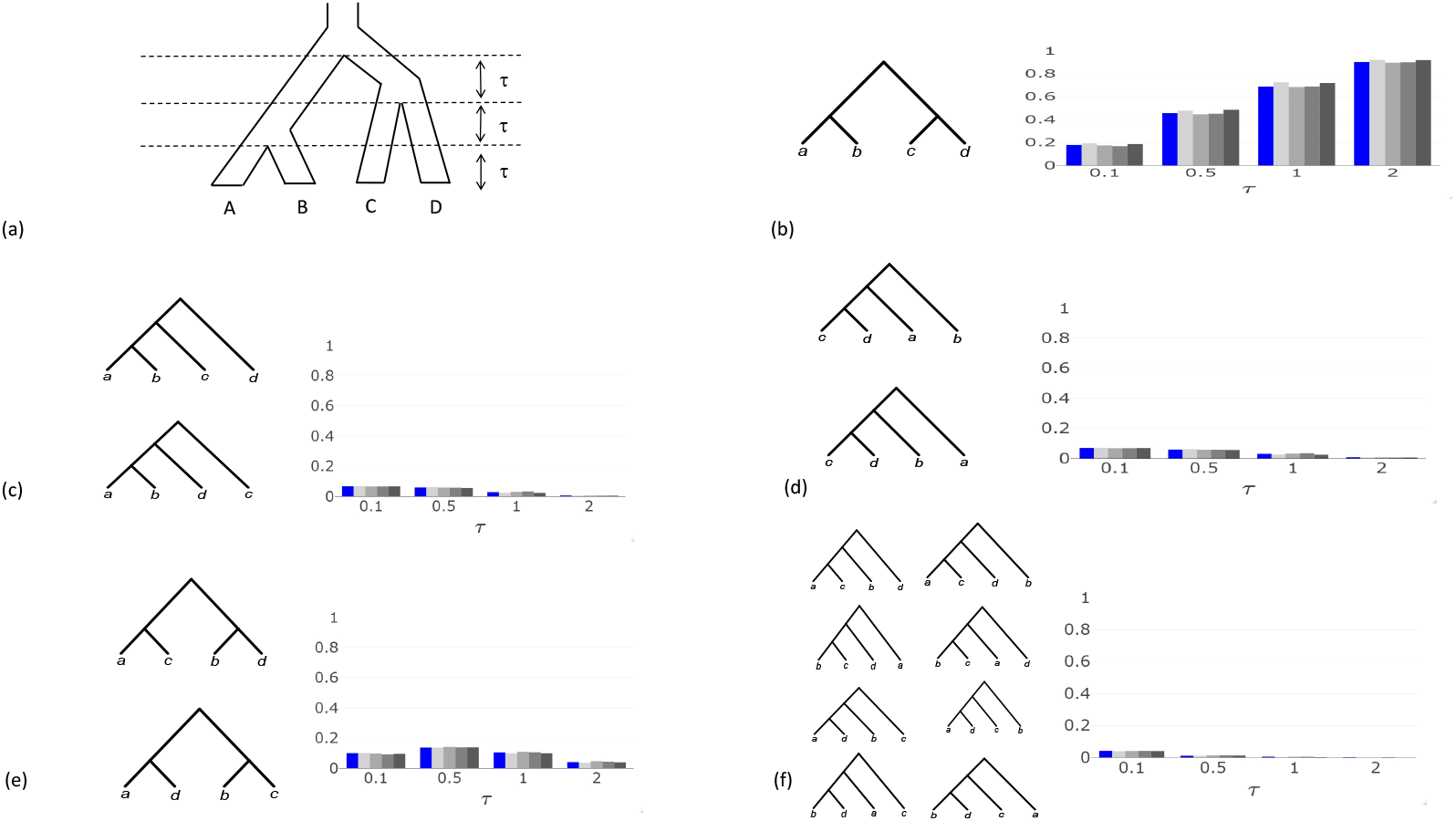
Gene tree probabilities for various choices of the selection and mutation parameters. (a) The species tree with internal branch lengths given by *τ*. Panels (b) - (f) give bar charts whether the height of the bars corresponds to the probability of the gene trees shown in that panel. Each cluster of bars gives probabilities for a different choice of *τ*. The blue bar shows probabilities for the neutral coalescent (*s* = 1). The other bars give probabilities for *s* = 1.0002, *p*_*m*_ = .0001 (light gray, left); *s* = 1.0002, *p*_*m*_ = .1 (medium-light gray, middle-left); *s* = 2, *p*_*m*_ = .0001 (medium-dark gray, middle-right); *s* = 2, *p*_*m*_ = .1 (dark gray, right).

Figure 4 shows the distribution of the time of the first coalescent event in the case of the each of the 6 possible such events when *s* = 1.0002 and *p*_*m*_ = .0001. Panel (a) shows the case when the species tree is the symmetric tree (see Figure 3(a) with *τ* = 1), and panel (b) shows the case when the species tree is asymmetric. The distributions show that the first coalescent time tends to be smaller under selection, but this difference is small. The code used to generate these simulations can be found at https://github.com/mwwascher/selection.

**Figure 4:**
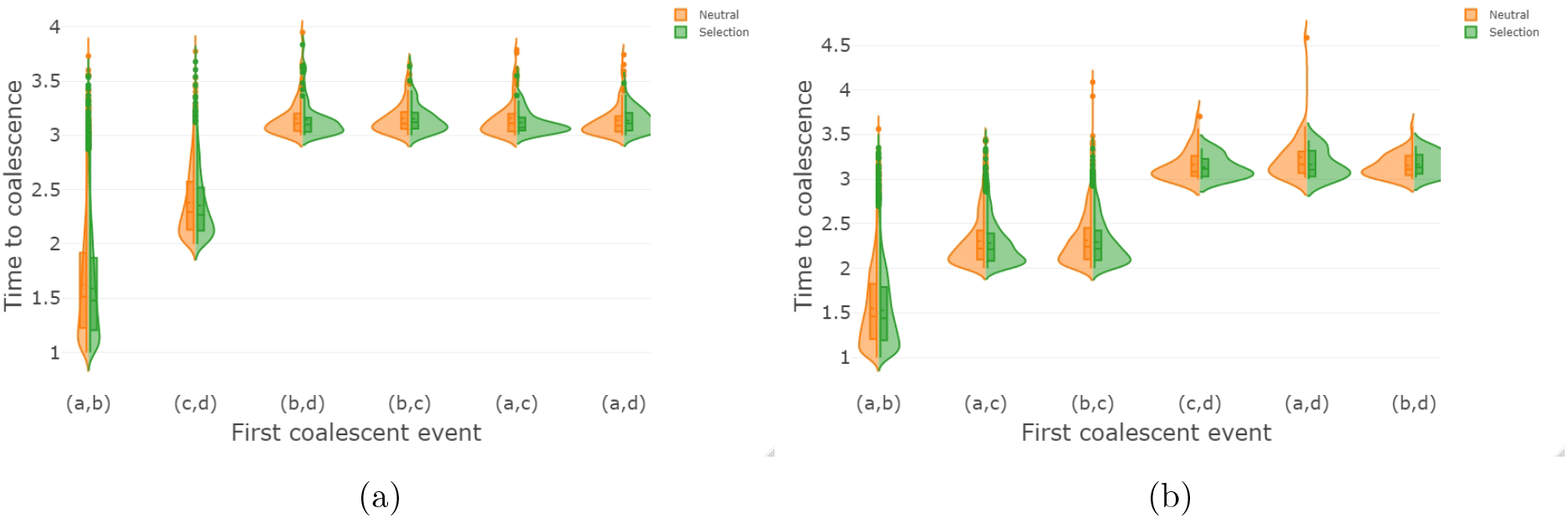
Distribution of the time to the first coalescent event when *s* = 1.0002, *p*_*m*_ = 0.0001, and *τ* = 1 for (a) the symmetric species tree in Figure 3(a) and (b) the asymmetric species tree in Figure x.

## 3 Discussion

We have derived an approximation for the rate of coalescence under a model with mutation and selection, and have demonstrated empirically that both the distribution of the number of type *A* alleles and the time to coalescence are well-approximated by our model. These results are then used to show that selection has only a moderate effect on both the coalescent times and the distribution of gene tree topologies. Here we discuss the approximation in terms of some theoretical heuristics, as well as broader implications for species-level phylogenomic inference.

### 3.1 Theoretical considerations for the approximation

Our simulations in Section 2.2.1 support that the approximation is adequate for studying coalescent-time distributions in many cases. However, we might still consider what the theory suggests about the approximation generally. Although we cannot rigorously derive the rate of convergence for the general case, we can use some heuristics to think about what the theory suggests in different scenarios.

The theory underlying our approximation is derived under the assumption that *s* and *p*_*m*_ are fixed while *N* → ∞. While in practice population sizes are not infinite, our approximation will apply to cases where the population size is reasonably large, and should not be used to make conclusions for populations that are small. In addition, the adequacy of our approximation depends on the particular values of *p*_*m*_ and *s*, as demonstrated by Figure 1 in Section 2.2.1. Below we discuss what the theory suggests about the approximation in four specific cases: for *p*_*m*_, we consider both large and small value of *p*_*m*_, where *p*_*m*_ is considered to be small when it is on the order 1*/N* ; for *s*, we consider large values to be those much greater than 1, while small value of *s* are those near 1.

#### Case 1: p_m_ large, s large

When both *p*_*m*_ and *s* are large, our approximation is good, as shown by our simulation results. However, this situation is unlikely to be common for empirical data.

#### Case 2: p_m_ large, s small

In this case, no matter the starting proportion of type *A* alleles, mutation overwhelms selection and we quickly reach *p* ≈ 1*/*2. Both the theoretical results and the simulations suggest that the approximation will hold well in this case. However, this case is also likely to be rare in empirical settings, since mutation rates are generally small.

#### Case 3: p_m_ small, s large

In this case, *p* is very close to 1 and the coalescent under selection will behave almost identically to the neutral coalescent. Our simulations confirm the theoretical prediction that provided that the initial value of *p*_*i*_ is not too close to 0, *p*_*i*_ quickly reaches *p* and the approximation holds well. Note, however, that if the initial proportion of *A* alleles is very close to 0, then *p*_*i*_ will stay close to 0 for a long time, until a critical mass of type *A* alleles accumulate through mutation, at which time *p*_*i*_ will jump quickly to *p*. However, since *p* is close to 1 in this case, and *p*_*i*_ close to 0 and *p*_*i*_ close to 1 both result in a coalescent process nearly identical to that of the neutral coalescent, the approximation will still hold well, regardless of the starting proportion of type *A* alleles.

We also note that this case provides some justification for analyses that ignore the effects of selection, in that ignoring selection implies that *p* = 1 or *p* = 0, which is close to what our approximation predicts. Thus, the distribution of coalescence times should be well-approximated by the neutral coalescent model when *s* is large and *p*_*m*_ is small.

#### Case 4: p_m_ small, s small

Our approximation does not fit as well in this case as for the others. In particular, our simulations show that as *p*_*m*_ and *s* − 1 become small compared to 1*/N*, what we observe empirically begins to differ from what the approximation predicts. Our theoretical arguments suggest that this is due to slow mixing of *h* in this case. However, our simulations suggest that even if the approximation does not hold perfectly, selection still appears to increase the rate of coalescence, which has important implications for species tree inference in the presence of selection.

### 3.2 Implications for species-level phylogenomic inference

We now turn our attention to inference and consider the consequences of using standard coalescent-based models for species tree inference that do accommodate selection when in fact selection is present. Our first observation is that in some cases, our results suggest the coalescent under selection behaves essentially identically to the neutral coalescent, in which case ignoring selection will not create any problems with inference. In particular, this is true when selection is strong and mutation rates are small, a scenario that may occur in practice. We note that this is also the case if mutation rates are much larger than *s* − 1, but this situation seems less likely to occur in empirical settings.

When *s*−1 and *p*_*m*_ are comparable in magnitude, our results suggest that rates of coalescence will be noticeably but not greatly different from those in the neutral coalescent. The most obvious result of this is that any estimates of coalescent and speciation times will be slightly biased upward. We note that this bias is not the product of a constant rescaling of time over the entire tree. Rather, the bias depends on the number of lineages at any given time, making it nontrivial to adjust for the bias rigorously, even if selection and mutation rates could be reliably estimated.

A striking conclusion of our work is that when primary interest is in estimating the species tree topology, the presence of selection can only help, even if it is ignored and the neutral coalescent is used, as has been predicted by others (e.g., Rannala and Yang (2003); Zhu and Yang (2012); Edwards (2009); Edwards et al. (2016)). This is because a higher rate of coalescence means that ILS is less likely, which results in a higher likelihood that the gene tree for a locus under selection will match the species tree. Thus, site patterns sampled from genes under selection are more likely to support the true species tree topology.

It is enticing to then believe that selection might be leveraged to improve estimates of the species tree topology. However, we feel this is not very promising for the following reason. The increased probability of seeing a gene tree that matches the species tree is small or practically none in many cases, such as the case of strong selection, and so unless selection and mutation rates can be estimated precisely, the small gains from utilizing loci under selection will likely be washed out by the uncertainty in the estimates of the selection and mutation rates. It is also not clear that a sufficient number of loci under selection will be available for inference in a wide range of empirical settings, a difficulty compounded by the fact that identification of loci under selection can be challenging.

Overall, our theory and simulation results agree with what has been previously observed concerning the impact of selection on species-level phylogenetic inference (e.g., Adam et al. (2018); Rannala and Yang (2003); Zhu and Yang (2012); Edwards (2009); Edwards et al. (2016)), but we provide a more concrete theoretical basis for these predictions. Our results have important implications for the practice of empirical phylogenetics when selection is thought to have played a role in the past history of the species under consideration.

## 4 Materials and Methods

### 4.1 Population dynamics

In this section, we derive some lemmas that follow from our definition of the Markov chain that tracks the portion of *A* alleles in the population for the Wright-Fisher model with selection, as defined in Section 2.1.1 and Lemma 2.1.

#### Lemma 4.1

*For the Markov chain* 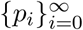 *in Lemma 2.1, let* 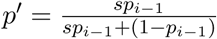. *Then*

1. 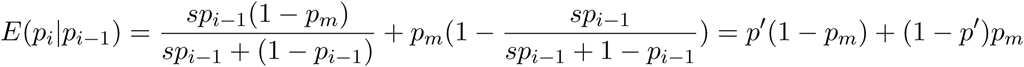
2. 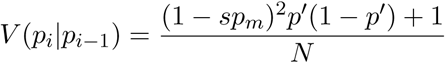

*Proof.*.

1. 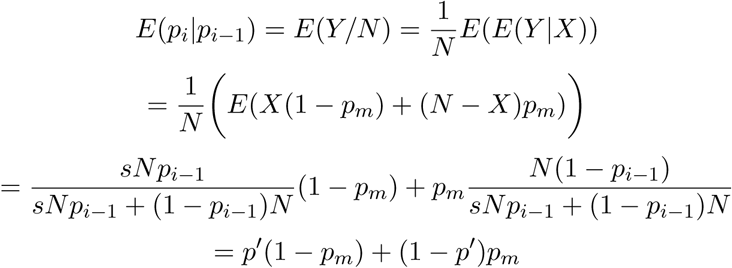
2. 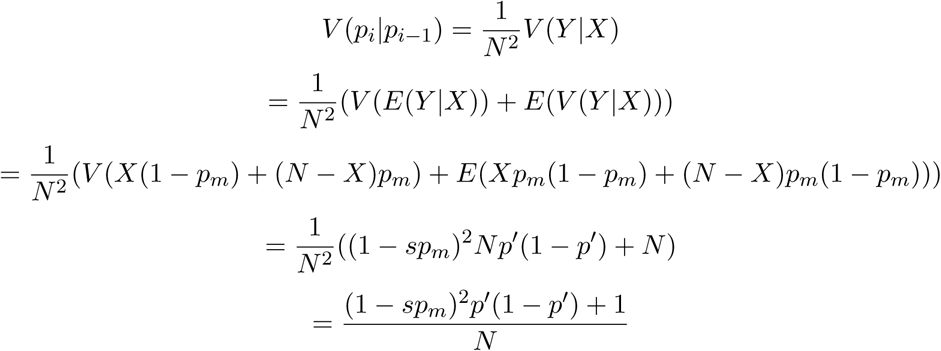

□

We next show that 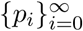 has a fixed point *p* such that *E*(*p*_*i*_|*p*_*i*−1_ = *p*) = *p*. Setting *p*_*i*_ = *E*(*p*_*i*+1_|*p*_*i*_) and solving for *p*_*i*_ gives

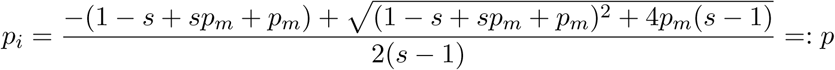

which is in [0, 1] when *s >* 1, *p*_*m*_ ∈ [0, 1].

It is also the case that *p*_*i*_ tends to move toward *p* and moves more toward *p* when it is further away. We can see this by looking at the derivatives of *E*(*p*_*i*_|*p*_*i*−1_).

#### Lemma 4.2

*The following hold:*

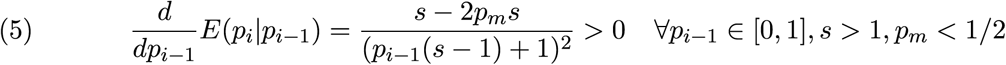

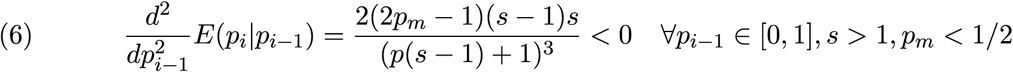

*Proof*. This can be shown using simple algebra. □

We next prove some lemmas that will be needed later.

#### Lemma 4.3

*For any p*_*i*_ ∈ [0, *p*), *E*(*p*_*i*+1_|*p*_*i*_) ∈ (*p*_*i*_, *p*) *and for any p*_*i*_ ∈ (*p*, 1], *E*(*p*_*i*+1_|*p*_*i*_) ∈ (*p, p*_*i*_).

*Proof*. Suppose *p*_*i*_ ∈ [0, *p*). Then it is sufficient to show that the graph of the function *f* (*p*_*i*_) = *E*(*p*_*i*+1_|*p*_*i*_) lies above the graph of the function *h*(*p*_*i*_) = *p*_*i*_ on the interval [0, *p*). It is easy to see that *E*(*p*_*i*+1_|*p*_*i*_ = 0) = *p*_*m*_ and *E*(*p*_*i*+1_|*p*_*i*_ = 1) = 1 − *p*_*m*_, and we know from the definition of *p* that *E*(*p*_*i*+1_|*p*_*i*_ = *p*) = *p*. Now suppose there exists *p*^∗^ ∈ (0, *p*) such that *E*(*p*_*i*+1_|*p*_*i*_ = *p*^∗^) ≤ *p*^∗^. Then the mean value theorem implies that

1. There exists a point in (0,*p\**) such that 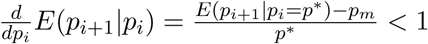.
2. There exists a point in (*p\**,1) such that 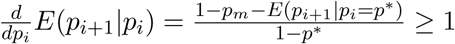.

However, this would create a contradiction, since Lemma 4.2 says that *E*(*p*_*i*+1_|*p*_*i*_) is an increasing concave function on [0, 1]. Thus, no such *p*^∗^ exists in [0, *p*) such that *E*(*p*_*i*+1_|*p*_*i*_ = *p*^∗^) ≤ *p*^∗^, and so for any *p*_*i*_ ∈ [0, *p*), *E*(*p*_*i*+1_|*p*_*i*_) ∈ (*p*_*i*_, *p*). An analogous argument can be used to show that for any *p*_*i*_ ∈ (*p*, 1], *E*(*p*_*i*+1_|*p*_*i*_) ∈ (*p, p*_*i*_).

#### Lemma 4.4

*The following hold:*

1. *For δ sufficiently small, E*(*p*_*i*+1_|*p*_*i*_) − *p*_*i*_ ≥ *E*(*p*_*i*+1_|*p*_*i*_ = *p* − *δ*) − (*p* − *δ*) *for all p*_*i*_ ≤ *p* − *δ*.
2. *For δ sufficiently small, E*(*p*_*i*+1_|*p*_*i*_) − *p*_*i*_ ≥ *E*(*p*_*i*+1_|*p*_*i*_ = *p* + *δ*) − (*p* + *δ*) *for all p*_*i*_ ≥ *p* + *δ*.
3. *For any E >* 0, ℙ (|*E*(*p*_*i*+1_|*p*_*i*_) − *p*_*i*+1_| *> ϵ*) ≤ 2 exp(−*N ϵ*^2^*/*6).

*Proof*. 1. holds because by lemma 4.2, *E*(*p*_*i*+1_|*p*_*i*_) − *p*_*i*_ is a smooth concave function that passes through (0, *p*_*m*_) and (*p*, 0). At some point in (0, *p*), *E*(*p*_*i*+1_|*p*_*i*_) − *p*_*i*_ must begin decreasing so it will reach 0 when *p*_*i*_ = *p*. Because it is concave, once this happens, it will not begin increasing again, so if *E*(*p*_*i*+1_|*p*_*i*_) − *p*_*i*_ is decreasing when *p*_*i*_ = *p* − *δ*, then 1. holds for that choice of *δ*.

2. holds for reasons analogous to 1.

3. is an application of a Hoeffding’s inequality for a binomial random variable. When going from the *i*th to the *i* + 1st generation, we can think of generating each offspring’s type by flipping a coin independently (since we choose a parent independently with replacement.) The probability of getting type A on this flip of course depends on *p*_*i*_, but given *p*_*i*_, *p*_*i*+1_ is a binomial random variable and so we can apply Hoeffding’s inequality which yields

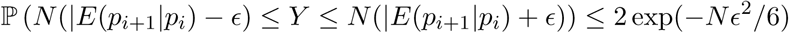

where 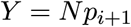 □

Rather than attempt to track *p*_*i*_ from generation to generation, we will show that under these dynamics *p*_*i*_ quickly approaches *p* and stays near *p* for a long time. So then asymptotically, we can examine the coalescent under the assumption that *p*_*i*_ = *p* for all generations.

### 4.2 Concentration of *{p*_*i*_*}* to *p*

Our goal for this section is to show that the Markov chain 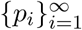 concentrates around *p* quickly and stays there for a long time when *N* is sufficiently large.

#### Lemma 4.5

*Fix ϵ >* 0 *and δ >* 0 *sufficien tly s mall as in Lemma 4.4, and let p*_0_ ∈ [0, 1] *\* (*p* − *δ, p* + *δ*). *Then with probability at least* 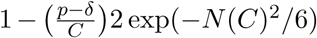 *there exists a constant C >* 0 *such that the process {p*_*i*_*} moves monotonically toward the interval* (*p* − *δ, p* + *δ*) *and reaches it in at most p/C steps*.

*Proof*. Without loss of generality, suppose that *p*_*i*_ *< p*−*δ*. Lemma 4.4 implies *E*(*p*_*i*+1_|*p*_*i*_)−*p*_*i*_ ≥ *h* where *h* = *E*(*p*_*i*+1_|*p*_*i*_ = *p* − *δ*) − (*p* − *δ*) *>* 0. Then if |*E*(*p*_*i*+1_|*p*_*i*_) − *p*_*i*+1_| ≤ *h/*2, we must have *p*_*i*+1_ − *p*_*i*_ ≥ *h/*2. At this point, either

1. *p*_*i*+1_ ≤ *p* − *δ*, in which case we can repeat the argument above to show that *p*_*i*+2_ − *p*_*i*+1_ ≥ *h/*2, or
2. *p*_*i*+1_ *> p* − *δ*. In this case, we must have *p*_*i*+1_ ∈ (*p* − *δ, p* + *δ*) because
  a. *E*(*p*_*i*+1_|*p*_*i*_) *< p* by Lemma 4.3.
  b. *E*(*p*_*i*+1_|*p*_*i*_ = *p* − *δ*) ∈ (*p* − *δ, p*) from Lemma 4.3 implies that *h < δ*.
  c. |*E*(*p*_*i*+1_|*p*_*i*_) − *p*_*i*+1_| ≤ *h/*2 implies that *p*_*i*+1_ ≤ *p* + *h/*2 *< p* + *δ*.

No matter where *p*_*i*_ starts in [0, *p* − *δ*], *p*_*i*_ will increase until it reaches the interval (*p* − *δ, p* + *δ*) by taking steps of at least size *h/*2, and so since *p*_*i*_ ≥ 0, it takes at most 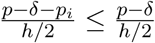 steps and this quantity does not depend on *N*.

Of course, for this to hold, we need |*E*(*p*_*i*+1_|*p*_*i*_) − *p*_*i*+1_| ≤ *h/*2 to hold for every step. Let *s* be the number of steps. Then using a union bound and applying Lemma 4.4 this probability is

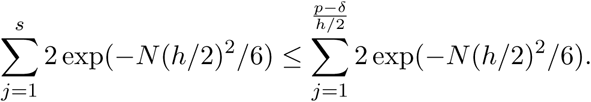

An analogous argument can be repeated for the case where *p*_*i*_ *> p* + *δ*. □

#### Theorem 4.6

*Fix ϵ >* 0 *and δ >* 0 *sufficiently small as in lemma 4.4*. *Let p*_0_ ∈ [0, 1] *and choose any k* ∈ ℕ. *Then there exists N* ^∗^ *such that for all N > N* ^∗^,

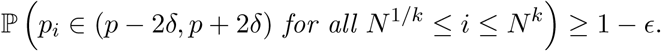

*Proof*. Note that

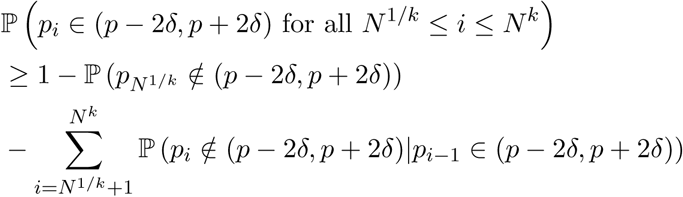

Lemma 4.5 implies that regardless of the value of *p*_0_, *p* _*i*_ will reach the interval (*p* − *δ, p* + *δ*) in at most 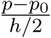 generations with probability at least 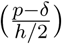 2 exp(−*N* (*h/*2)^2^*/*6), so we must first choose *N* ^∗^ such that 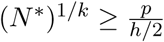, in which case

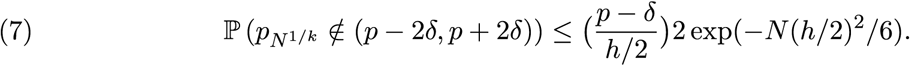

Suppose now that we observe *p*_*i*_ ∈ (*p* − *δ, p* + *δ*) for some 2 ≤ *i* ≤ *N* ^1*/k*^. Supposing that |*E*(*p*_*i*+1_|*p*_*i*_) − *p*_*i*+1_| ≤ *h/*2 for the value of *h* for the next step in Lemma 4.5, then either

1. *p*_*i*+1_ ∈ (*p* − *δ, p* + *δ*) in which case we simply let the process take another step, or
2. *p*_*i*+1_ ∉ (*p* − *δ, p* + *δ*). In this case, we must still have *p*_*i*+1_ ∈ (*p* − 2*δ, p* + 2*δ*) because lemma 4.3 implies that *E*(*p*_*i*+1_|*p*_*i*_) ∈ (*p* − *δ, p* + *δ*), and together with |*E*(*p*_*i*+1_|*p*_*i*_) − *p*_*i*+1_| ≤ *h/*2 *< δ* implies that *p*_*i*+1_ ∈ (*p* − 2*δ, p* + 2*δ*). In this case we apply Lemma 4.5 again to show that *{p*_*i*_*}* quickly reaches (*p* − *δ, p* + *δ*) again with high probability.

In order for *{p*_*i*_*}* to leave (*p* − 2*δ, p* + 2*δ*), we must observe a generation where either

1. |*E*(*p*_*i*+1_|*p*_*i*_) − *p*_*i*+1_| *> h/*2, or
2. Lemma 4.5 fails, which also happens when |*E*(*p*_*i*+1_|*p*_*i*_) − *p*_*i*+1_| *> h/*2.

Since we need |*E*(*p*_*i*+1_|*p*_*i*_) − *p*_*i*+1_| *> h/*2 to be satisfied for at most *N* ^*k*^ generations, the union bound on the probability of failure yields

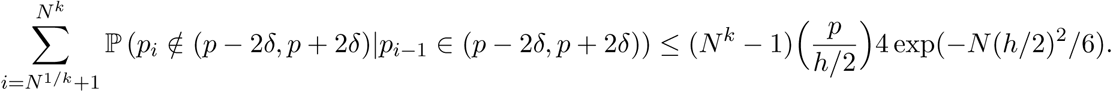

Combining this with 7, we conclude

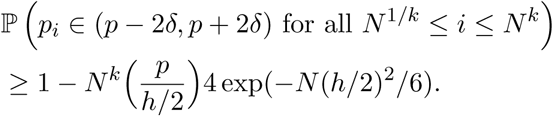

Thus also choosing *N* ^∗^ large enough so that 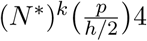 exp(−*N* (*h/*2)^2^*/*6) *< ϵ* completes the proof. □

### 4.3 Deriving the coalescent under selection

Intuitively, it is not hard to see that regardless of the value of *p*_*i*_, the probability of seeing a least one coalescent event from generation *i* to generation *i* − 1 is at least *O*(1*/N*). Since the population has *N* individuals in generation *i*, at least one of the number of individuals with type *A* parents in generation *i* − 1 or the number of individuals with type *a* parents in generation *i* − 1 is *O*(*N*). If we choose the larger of the two, then it has size *O*(*N*), and the probability of observing a coalescent event among only these individuals is *O*(1*/N*). This can be seen by using the results from the neutral Wright-Fisher model, since individuals of the same type are exchangeable.

As described above, it is reasonable to assume that asymptotically the coalescent process behaves as if *p*_*i*_ = *p* for all *i*. Here we give details of the approximation of the waiting time until the next coalescent event with the exponential distribution with parameter *λ* as given in Equation 4, when time is appropriately scaled.

Equation 3 above gives the probability of one coalescent event in the immediately previous generation. We assume that the probability of multiple coalescent events is *o*(1*/N*). We would like to now treat the number of generations to the coalescent event as a geometric random variable. However, unlike in the standard coalescent, the probability of success for our geometric random variable varies from generation to generation because it depends on the value of *h*.

We can address this by considering the Markov chain 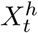 that tracks the value of *h* in *{*0, …, *M }* in each generation. Since all transitions have positive probability, the chain is ergodic and has a stationary distribution *π* that can be easily derived by solving the detailed balance equations for small to moderate *M*. In practice, we conceive of *M << N* (usually *M* = *O*(1),) and in this case the Markov chain mixes quickly compared to *N* since the transition probabilities do not depend on *N* in any way, but rather on *M, p, s, p*_*m*_. We can show this as follows:

Choose the initial state of 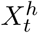 arbitrarily, and let 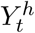 be another chain on the same space with the same transition matrix as 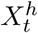, but choose the initial state of 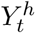 according to *π*. We will couple 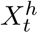 and 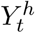 as follows:

1. If 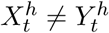, choose the transitions for 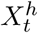 and 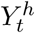 independently according the transition matrix.
2. If 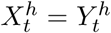, choose a single transition according to the transition matrix and apply it to both chains.

Note that in this coupling, once we have 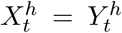, this persists forever. Additionally, the distribution of 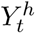 is at all times the stationary distribution, so the first time we observe 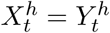, the chain has mixed in the sense that if *τ* is this time then 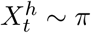 for all *t* ≥ *τ*.

Now suppose *p*_*ij*_ are the entries of the transition matrix. In any step *t* with 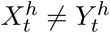, there is always some probability that we will see 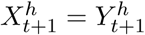 in the next step, since any state can be reached in one step from any other state, and this probability must be at least min_*i,j*_(*p*_*ij*_)^2^ *>* 0. So the time it takes the chain to mix is dominated above by a geometric random variable with success probability min_*i,j*_(*p*_*ij*_)^2^, which may depend on *M, p, s, p*_*m*_ but not *N*. Thus if we take some slowly growing function of *N* such as log *N*, then

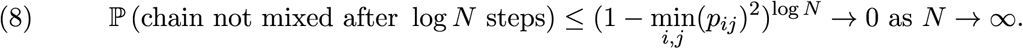

Since *N* is large compared to *M*, the proportion of time spent at each value of *h* over the course of *O*(*N*) generations will be about *π*(*h*). Additionally, while we use *π*(*h*) here for simplicity of notation, *π*(*h*) is in fact a function of *s, p*_*m*_, and *M* in addition to *h* that can easily be computed for specific values.

Now counting time in units of *N* generations, let *T* ^*N*^ be the time until the coalscent event from *M* lineages to *M* − 1 lineages counted in time units *t* of *N* generations. Then

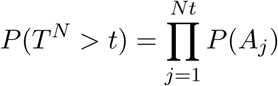

where *A*_*j*_ is the event that there is not a coalescent event in generation *j*. This is the same as

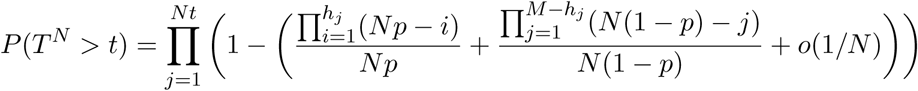

where *h*_*j*_ is the value of *h* in generation *j*. Let *π*(*h*) + *ϵ*_*N*_ (*h*) be the proportion of time *h*_*j*_ spend in each state *h* = 1, …, *M*. Then

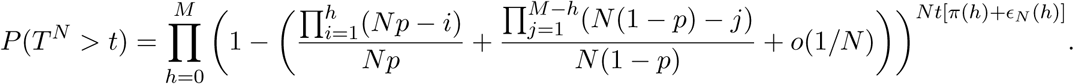

We can rewrite this as

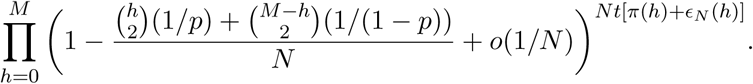

Recall that one way to interpret *π*(*h*) is that when viewed on a timescale large compared to its mixing time, *h*_*j*_ will spend a proportion of time approaching *π*(*h*) in state *h* for each *h* = 1, …, *M*. Equation 8 shows that with high probability *h*_*j*_ mixes after log *N* time, and so in our asymptotic setting, *N* is such a timescale, since *N/* log *N* → ∞ as *N* → ∞. Accordingly we may fix *ϵ >* 0 and choose *N* ^∗^ large such that with high probability | *ϵ*_*N*_ (*h*)| *< E* for *h* = 1, …, *M* and all *N > N\**. In this case we observe

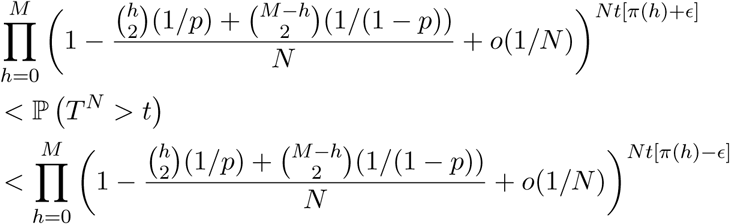

and as *N* → ∞, the upper and lower bounds converge to

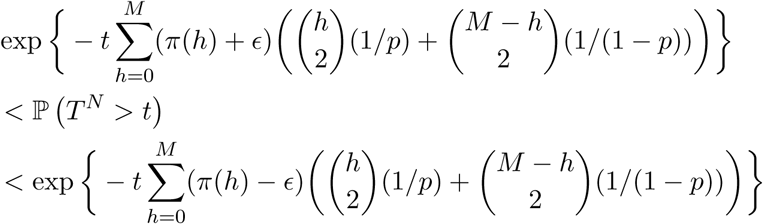

where *ϵ >* 0 can be chosen to be as small as desired in the previous step. We thus conclude that the limiting distribution for the coalescent time is an exponential distribution with rate parameter

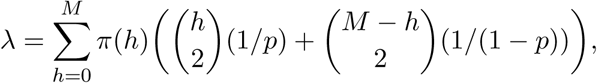

as given in Equation 4.

As in Kingman’s coalescent, the waiting time until the next coalescent event is exponentially distributed, but this distribution now depends on the selection advantage *s* and the mutation rate *p*_*m*_ in addition to the number of lineages *M*. Note that *π*(*h*) is in fact a function of *M, s*, and *p*_*m*_ in addition to *h*.

### 4.4 Description of the algorithm used to simulate coalescent times

We simulated the coalescent times shown in Figure 4 using Julia with the following algorithm, which we describe briefly below. Our code can be found at https://github.com/mwwascher/selection.

The steps to a simulate a coalescent time are as follows:

1. Choose parameters: choose the desired values of selection advantage *s*, mutation rate *p*_*m*_, and population size *N*. Set *j* = 2*N*
2. Simulate 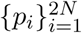: set *p*_1_ = .5. For *i* = 2, …, 2*N*, draw *Y* |*p*_*i*−1_ as described in Lemma 2.1 and set *p*_*i*_ = *Y/N*.
3. Initialize the lineages: Draw *M* lineages and their types uniformly at random from the *N* individuals in the population at time 2*N*.
4. Determine parent types: For each of the *M* lineages, determine that lineage’s parent type given the current value of *p*_*j*_ using the probabilities in equation 2. Let *h* be the number of lineages with type *A* parents.
5. Check for a coalescent event: Flip a coin with probability of success given by equation 3 to see if a coalescent event occurs in the parent generation. If a coalescent event occurs or *j* = 1, stop and output the coalescent time (in coalescent units) of *j/N*. Otherwise, set *j* = *j* − 1 and go to step 4.

The algorithm ends when a coalescent event has been observed or 2 coalescent units of time have passed. These steps can be repeated to sample multiple coalescent times. For our simulations, we used *N* = 500000 for the case when *M* = 6 and *N* = 200000 for the case when *M* = 4 and simulated 1000 coalescent times for each combination of *s* ∈ *{*2, 1.0002*}, p*_*m*_ ∈ *{*0.1, 0.0001*}*

## References

Adam, R. H., D. R. Schield, D. C. Card, and T. A. Castoe. 2018. Assessing the impacts of positive selection on coalescent-based species tree estimation and species delimitation. Systematic Biology 67:1076–1090.

Barton, N. and A. Etheridge. 2019. Mathematical Models in Population Genetics chap. 4, Pages 115–20. John Wiley & Sons, Ltd.

Castoe, T., A. P. J. de Koning, H.-M. Kim, W. Gu, B. P. Noonan, G. N. G, Z. J. Jiang, C. L. Parkinson, and D. D. Pollock. 2009. Evidence for an ancient adaptive episode of convergent molecular evolutio. Proceedings of the National Academy of Sciences 106:8986–8991.

Corbett-Detig, R., D. L. Hartl, and T. Sackton. 2015. Natural selection constrains neutral diversity across a wide range of species. PLoS Biology 13.

Edwards, S. C. 2009. Is a new and general theory of molecular systematics emerging? Evolution 63:1–19.

Edwards, S. V., Z. Xi, A. Janke, B. C. Faircloth, J. E. McCormack, T. C. Glenn, B. Zhong, S. Wu, E. M. Lemmon, A. R. Lemmon, A. D. Leaché, L. Liu, and C. C. Davis. 2016. Implementing and testing the multispecies coalescent model: a valuable paradigm for phylogenomics. Molecular Phylogenetics and Evolution 94:447–462.

Flouris, T., X. Jiao, B. Rannala, and Z. Yang. 2018. Species tree inference with bpp using genomic sequences and the multispecies coalescent. Molecular Biology and Evolution 35:2585–2593.

Hahn, M. 2008. Toward a selection theory of molecular evolution. Evolution 62:255–265.

Kaplan, N. L., T. Darden, and R. R. Hudson. 1988. The coalescent process in models with selection. Genetics 120:819–829.

Kimura, M. 1955. Random genetic drift in multi-allelic locus. Evolution 9:419–435.

McVicker, G., D. Gordon, C. Davis, and P. Green. 2009. Widespread genomic signatures of natural selection in hominid evolution. PLoS Genetics 5:e1000471.

Rannala, B. and Z. Yang. 2003. Likelihood and Bayes estimation of ancestral population sizes in hominoids using data from multiple loci. Genetics 164:1645–1656.

Rannala, B. and Z. Yang. 2017. Efficient Bayesian species tree inference under the multispecies coalescent. Systematic Biology 66:823–842.

Scally, A., J. Y. Dutheil, L. Hillier, G. E. Jordan, I. Goodhead, J. Herrero, A. Hobolth, T. Lappalainen, T. Mailund, T. Marques-Bonet, S. McCarthy, S. H. Montgomery, P. Schwalie, Y. Tang, M. Ward, Y. Xue, B. Yngvadottir, C. Alkan, L. Andersen, Q. Ayub, E. Ball, K. Beal, B. Bradley, Y. Chen, C. Clee, S. Fitzgerald, T. Graves, Y. Gu, P. Heath, A. Heger, E. Karakoc, A. Kolb-Kokocinski, G. Laird, G. Lunter, S. Meader, M. Mort, J. Mullikin, K. Munch, T. O’Connor, A. Phillips, J. Prado-Martinez, A. Rogers, S. Sajjadian, D. Schmidt, K. Shaw, J. Simpson, P. Stenson, D. Turner, L. Vigilant, A. Vilella, W. Whitener, B. Zhu, D. N. Cooper, P. de Jong, E. Dermitzakis, E. E. Eichler, P. Flicek, N. Goldman, N. Mundy, Z. Ning, D. Odom, C. Ponting, M. Quail, O. Ryder, S. Searle, W. Warren, R. Wilson, M. Schierup, J. Rogers, C. Tyler-Smith, and R. Durbin. 2012. Insights into hominid evolution from the gorilla genome sequence. Nature 483:169–175.

Siepel, A. 2009. Phylogenomics of primates and their ancestral populations. Genome Research 19:1929–1941.

Silva, D. N., S. Duplessis, P. Talhinhas, H. Azinheira, O. S. Paulo, and D. Batista. 2015. Genomic patterns of positive selection at the origin of rust fungi. PLoS ONE 10:e0143959.

Springer, M. and J. Gatesy. 2016. The gene tree delusion. Molecular Phylogenetics and Evolution 94:1–33.

Svirezhev, Y. M. and V. P. Passekov. 1990. Diffusion Models of Population Genetics Pages 239–268. Springer Netherlands, Dordrecht.

Tong, K. J., S. Duchêne, N. Lo, and S. Y. W. Ho. 2017. The impacts of drift and selection on genomic evolution in insects. PeerJ 5:e3241.

Wakeley, J. 2009. Coalescent Theory: An Introduction. Roberts and Company.

Yang, Z. and B. Rannala. 2010. Bayesian species delimitation using multilocus sequence data. Proceedings of the National Academy of Sciences 107:9264–9269.

Zhang, C., D. X. Zhang, T. Zhu, and Z. Yang. 2011. Evaluation of a bayesian coalescent method of species delimitation. Systematic Biology 60:747–761.

Zhu, T. and Z. Yang. 2012. Maximum likelihood implementation of an isolation-with-migration model with three species for testing speciation with gene flow. Molecular Biology and Evolution 29:3131–3142.

